# Dexamethasone promotes breast cancer stem cells in obese and not lean mice

**DOI:** 10.1101/2021.06.18.449008

**Authors:** Stephanie Annett, Orla Willis Fox, Damir Vareslija, Tracy Robson

## Abstract

Obesity is highly prevalent in breast cancer patients and it is associated with increased recurrence and breast cancer specific mortality. Glucocorticoid (GC) use, in addition to obesity is associated with promoting breast cancer metastasis through activation of stemness-related pathways. Therefore, we utilised the synergetic allograft E0771 breast cancer model to investigate if treatment with GCs had differential effects on promoting cancer stem cells in lean and diet-induced obese mice. Indeed, both lean mice treated with dexamethasone and obese mice with no treatment had no effect on the *ex vivo* colony forming ability, mammosphere formation or ALDH bright subpopulation. However, treatment of obese mice with dexamethasone resulted in a significant increase in *ex vivo* colony formation, mammosphere formation, ALDH bright subpopulation and expression of pluripotency transcription factors. GC transcriptionally regulated genes were not altered in the dexamethasone treated groups compared to treatment controls. In summary, these results provide initial evidence that obesity presents a higher risk of GC induced cancer stemness via non-genomic GC signalling which is of potential translational significance.

## Introduction

Steroids are routinely used in patients with solid tumours to manage tumour and treatment related symptoms. However, a meta-analysis of >80,000 patients found that use of steroids in solid tumours was associated with reduced overall and progression free survival [1]. One explanation for this is the observation is that glucocorticoids (GCs) promote a stem cell phenotype [2–4]. In breast cancer, plasma levels of cortisol (an endogenous GC) increase during breast cancer progression, inducing activation of the glucocorticoid receptor (GR) resulting in increased metastatic colonisation [5]. Worryingly, treatment with dexamethasone (a clinically used GC drug) also activated the GR causing increased metastatic burden and decreased survival [5]. Obesity affects more than half a billion adults worldwide and it is a well-known risk factor for breast cancer and an indicator of poorer prognosis [6,7]. Indeed, obese breast cancer patients have an increased relative risk of recurrence of 40% to 50%, however, the underlying biology behind the link between obesity and breast cancer progression remains unclear [8]. Obesity causes chronic inflammation in the adipose tissue and elevated levels of cortisol in both the local adipose tissue and the systemic circulation [9,10]. Given the role of GCs in breast cancer metastasis, we hypothesised that obesity may increase the risk of GC-induced breast cancer reoccurrence by further promoting cancer stem cells.

## Methods

Female 8 week old C57BL/6N mice (Charles Rivers, UK) were fed normal chow diet (Lab Supply, Advanced Protocol PicoLab Verified 75) or high fat diet (Datesand, Mouse Diet High Fat Pellets) for 12 weeks. E0771 cells (CH3 BioSystems, USA) where authenticated by short tandem repeat profiling, screened for pathogens by the manufacturer and maintained in DMEM (Sigma, Ireland) with 10% foetal bovine serum (Sigma, Ireland) and 20 mM HEPES (Invitrogen, Ireland) at 37 C in an atmosphere of 95% air/5% CO_2_. 1 x 10^6^ E0771 murine breast cancer cells were resuspended in 100 μl Matrigel:PBS (1:1) and injected into the 4th mammary fat pad. Mice were randomly assigned to treatment groups (Fig. 1A; n>5 mice/group). *In vivo* dexamethasone treatment (water soluble dexamethasone; D2915, Sigma) was commenced when tumours were ∼150 mm^3^ at a clinically relevant dose of 0.1 mg/kg in drinking water for 5 out of 7 consecutive days for a total of three weeks [5]. Tumour volumes were measured and calculated three times weekly and excised when the tumour reached a mean diameter of 12 mm. For all *in vivo* experiments, mice were housed in individually ventilated cages according to EU Directive 2010/63 at constant temperature and humidity with 12-h light/dark cycle and fed standard chow. The welfare of all the mice was monitored daily and health screening carried out regularly as per the policy of licensed establishment. Mice were euthanised by using exposure to CO_2_. No adverse events were noted for *in vivo* experiments. The experimental protocols were compliant with the ARRIVE guidelines and Individual License Number I181 under the Project License Number P045.Tumours were dissociated and *ex vivo* analysis of mammospheres, colony forming ability and aldehyde dehydrogenase (ALDH) was determined as previously described [11–13]. RNA was isolated using the RNAeasy kit (Qiagen) and gene expression quantified by real-time PCR with primers listed in Supplementary Table 1.

**Figure 1.**
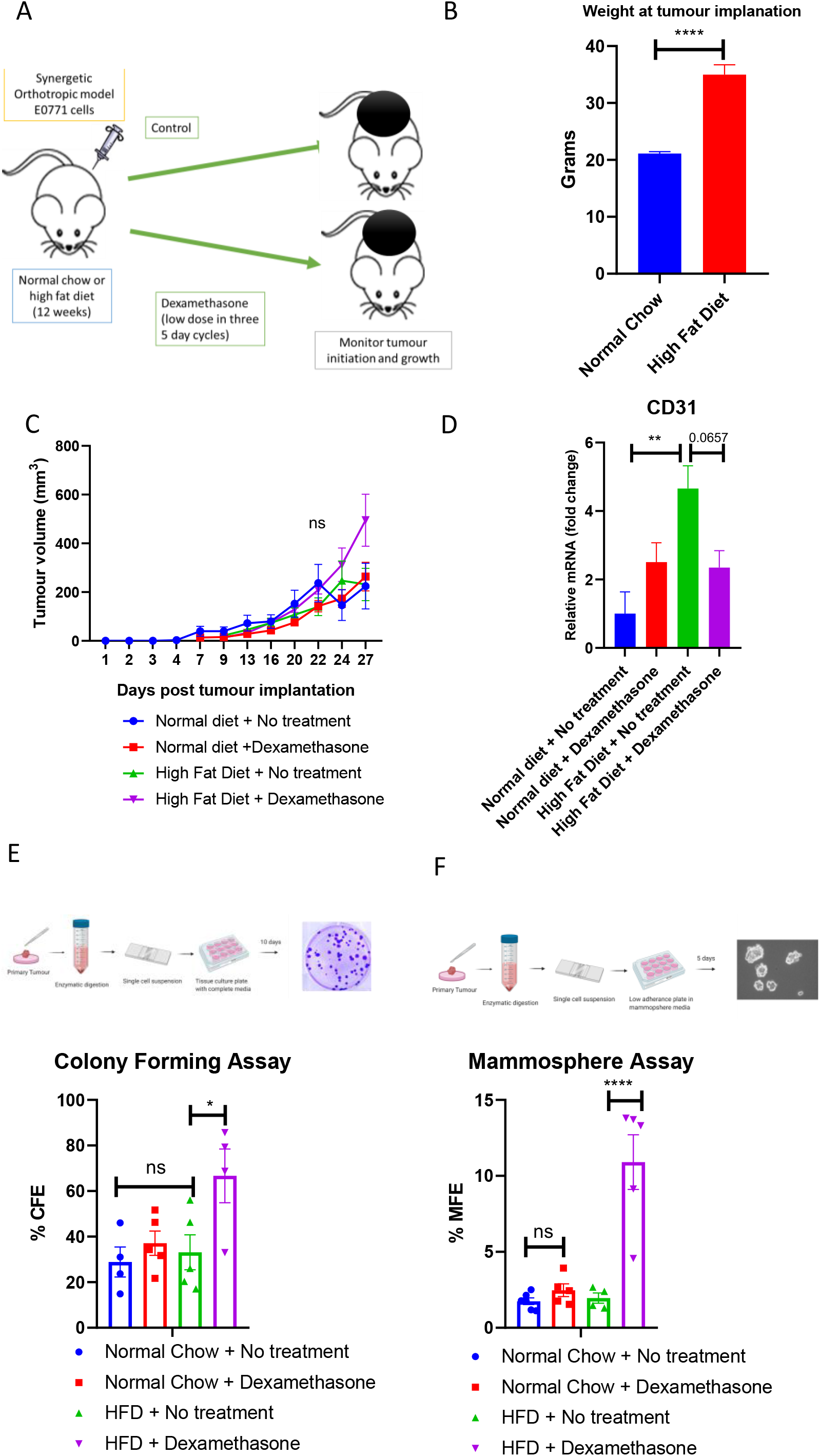
High fat diet and dexamethasone does not alter tumour growth but promotes *ex vivo* mammosphere and colony formation. **(A)** Overview of experimental design. Female C57BL/6N mice were fed a normal chow diet or high fat diet (HFD) for 12 weeks. E0771 murine tumour cells were implanted into the mammary fat pad and mice were randomly allocated no treatment or dexamethasone (0.1 mg/kg) in the drinking water for 5 out of 7 days for 3 weeks. Tumours were excised for *ex vivo* analysis. **(B)** Weight of mice following 12 weeks of normal chow diet or HFD. **(C)** Tumour growth rate of mice in the following treatment groups; normal chow diet and no treatment (n=10), normal chow diet and dexamethasone (n=7), HFD and no treatment (n=6), HFD and dexamethasone (n=5). **(D)** Expression of mRNA *Cd31* within excised E0771 tumours following normal diet or HFD and treatment with dexamethasone. **(E)** *Ex vivo* colony forming ability and **(F)** Mammosphere formation following different diets or treatment with dexamethasone. Data points are mean ± SEM. *n* ≥ 3. ^*^*P* < 0.05; ^**^*P* < 0.01 (one-way ANOVA or two-tailed Student *t*-test).

## Results

Female mice fed a high fat diet (HFD) for 12 weeks were significantly heavier than mice fed a normal diet (Fig. 1B; p<0.0001). Contrary to previous reports of diet induced obesity promoting E0771 tumour growth [14,15] we found no difference in the growth rate between any of the groups; normal diet and no treatment (n=10), normal diet and dexamethasone (n=7), HFD and no treatment (n=6), HFD and dexamethasone (n=5) (Fig. 1C). GCs were previously reported to have anti-angiogenic activity in solid tumours [16] and therefore we measured the gene expression of the endothelial marker *Cd31* (also known as *Pecam-1*). There was no difference in *Cd31* gene expression between mice on a normal diet treated with dexamethasone (Fig 1D). However, mice fed a HFD had a significant increased expression (Fig. 1D, p=0.0073), in line with previous reports of obesity inducing angiogenesis in breast cancer [15], and this was partially inhibited by dexamethasone treatment although it did not reach significance (Fig. 1D, p=0.0657). *Ex vivo* analysis of the dissociated tumour cells found there was no increase in colony forming ability in the mice on a normal diet treated with dexamethasone or mice fed a HFD (Fig. 1E). However, there was a significant increase in colony forming ability, indicative of enhanced clonogencity, in the mice fed a HFD and treated with dexamethasone (Fig. 1E, p=0.0432). In addition, mammosphere forming efficiency was analysed in the dissociated tumour cells. Similarly, there was no increase in mammospheres in mice on a normal diet treated with dexamethasone or those on a HFD (Fig. 1F). However, there was an approximately 10-fold increase in mammospheres in mice on a HFD treated with dexamethasone, compared to the other treatment groups, indicative of enhanced stemness (Fig. 1F; p<0.0001). Together these results indicate that tumour cells from obese mice treated with dexamethasone have increased self-renewal activity and cell survival compared to mice on a normal diet treated with dexamethasone. ALDH has been extensively studied in breast cancer as a marker for cancer stem cells. In addition, it is associated with chemo resistance and poor survival in patients [17]. Mice on a normal diet, or a HFD, did not have an increased ALDH^+^ bright population (Fig. 2A). However, in line with previous results mice fed a HFD and treated with dexamethasone had an increased ALDH^+^ bright subpopulation compared to the other treatment groups (Fig. 2A; p=0.0302). Next, we measured the pluripotency transcription factors *Nanog, Oct4* (also known as *Pou5F1*) and *Sox2. Oct4* expression was not detectable in the tumours of mice from any treatment group. *Sox2* gene expression was induced ∼ 6-fold by both dexamethasone in the mice fed a normal diet and in the mice fed a HFD and this increased to 10-fold in mice fed a HFD and treated with dexamethasone (Fig. 2B; normal diet vs normal diet + dexamethasone p=0.0003; normal diet vs HFD p=0.0003; HFD vs HFD + dexamethasone p=0.0025). Furthermore, HFD mice treated with dexamethasone also had a ∼ 10-fold increase in *Nanog* expression, compared to the other treatment groups (Fig. 2B; p<0.0001). It has been reported that GCs increase stemness and metastasis through activation of the non-canonical Wnt signaling axis [4,5] and therefore we investigated if *Ror1*, and its ligand *Wnt5a*, was also activated in our tumours. Treatment with dexamethasone on a normal diet increased expression of *Wnt5A* (Fig. 2C; p=0.0506), but this was not increased by HFD (Fig. 2C). *Ror1* expression was not altered in any of the treatment groups (Fig. 2C). In addition, genes classically regulated by GCs, *Nrcl1* and *Fkbp51* [18], were not induced in the tumours of any group (Fig. 2D). Together, this indicates that dexamethasone promotes a cancer stem cell phenotype in obesity through upregulation of pluripotency transcription factor via non-classical GC signaling.

**Figure 2.**
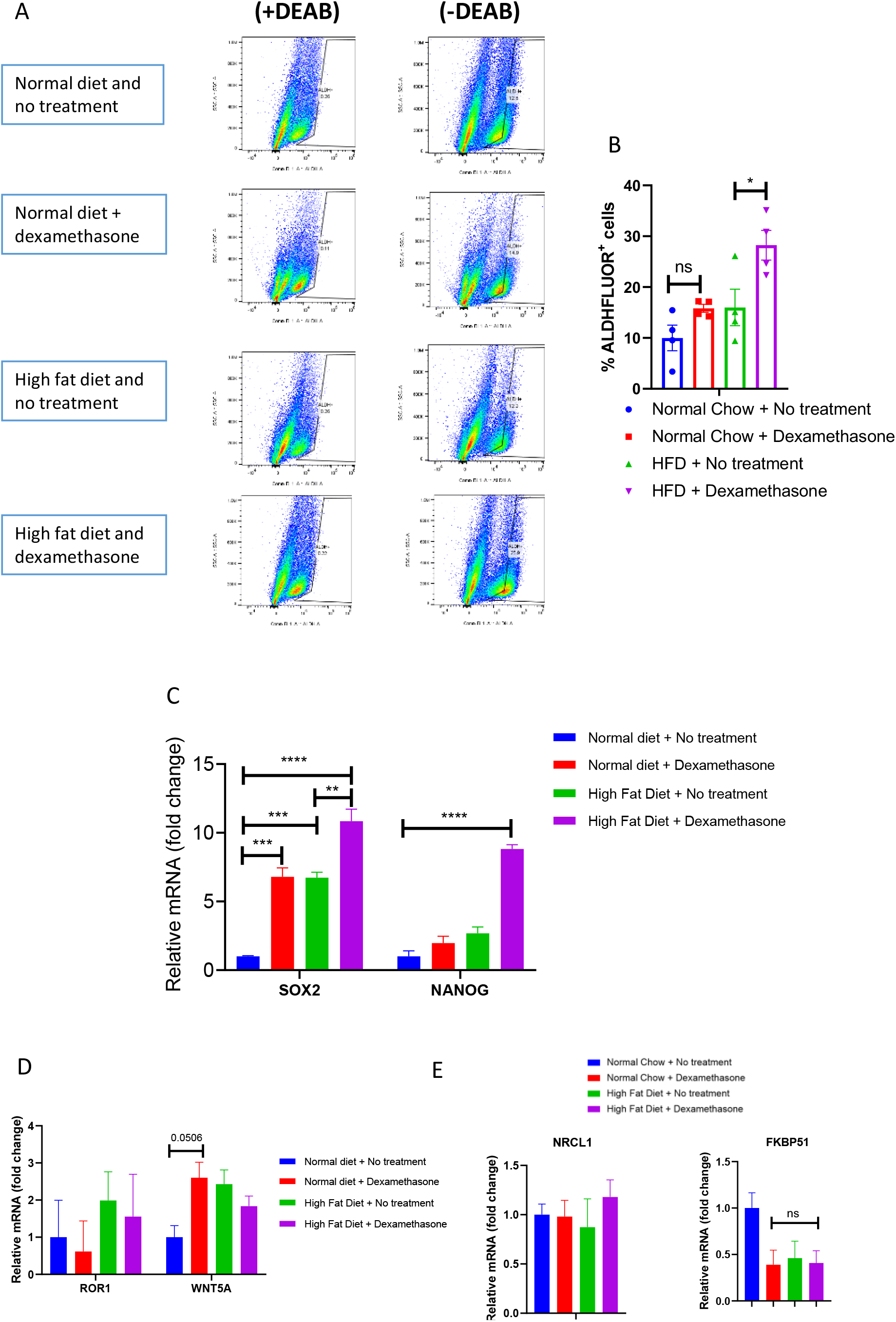
High fat diet and dexamethasone promotes an ALDH bright subpopulation and pluripotency transcription factors. **A)** *Ex vivo* analysis of an ALDH bright subpopulation of cells in excised E0771 tumours from mice fed either a normal or high fat diet and/or treated with dexamethasone (0.1 mg/kg) in the drinking water for 5 out of 7 days for 3 weeks. Expression of mRNA **(B)** *Sox2* and *Nanog*, **(C)** *Ror1* and *Wnt5a* and **(D)** *Nr3cl1* and *Fkbp51* from excised E0771 tumours following normal diet or HFD and treatment with dexamethasone. Data points are mean ± SEM. *n* ≥ 3. ^*^*P* < 0.05; ^**^*P* < 0.01 (one-way ANOVA or two-tailed Student *t*-test). SSC – side scatter

## Discussion

Adjuvant GC drugs (such as dexamethasone) are routinely used during the treatment of cancer to mitigate the undesirable effects of chemotherapy such as nausea and fatigue and to stimulate appetite [19]. In addition, they reduce hypersensitivity reactions to some chemotherapy agents and enhance tolerability to allow higher chemotherapy doses and more frequent cycles [19]. However, studies have revealed that GCs can promote cancer progression, nevertheless, the literature remains conflicting, and it is not clear how GCs either promote or inhibit tumour progression in different cancer types and by different mechanisms. Here, for the first time, we describe that obesity also has a profound effect on the ability of dexamethasone to promote the cancer stem cell phenotype, that does not occur in lean mice. Interestingly, overall, we found that obesity or dexamethasone alone had a limited effect on promoting cancer stem cells (Fig. 1F, Fig. 2A). This is contrast to previous reports demonstrating that both obesity and dexamethasone alone promotes metastasis via stem cell related signaling [3,4,14,20–23]. In our study, we found that dexamethasone promoted expression of the pluripotency transcription factor *Sox2* in both normal diet and obese mice (Fig. 2B), potentially indicating that a longer treatment time may promote stemness in lean mice. On the contrary, several studies have shown that dexamethasone decreases *Sox2* and thereby inhibits cancer stem cells [24,25]. In addition, similar to our study, dexamethasone did not promote *Ror1* expression in pancreatic cells [24] (Fig. 2C). Together these results indicate that GCs are a regulator of *Sox2* expression, however, its activation or inhibition is dependent upon the cell type and/or microenvironment. Indeed, a recent preprint on BioRxiv demonstrates, through single cell transcriptomics, that following acute GC exposure, cerebral organoids elicit differential effects depending on cell type [26]. Similar experiments utilizing single cell sequencing will aid our understanding of the cell type specific effects of GCs within the tumour microenvironment. Patients living with obesity make up a substantial proportion of individuals with breast cancer, however, they may not benefit equitably from established therapies. It is well-described that there is a reduced efficacy of cancer treatment among obese patients, particularly for chemotherapy in which the dose is often based on ideal body weight rather than actual body weight because of toxicity concerns [27]. We now provide initial evidence that GCs may have an altered mechanism of action in obesity resulting in expansion of breast cancer stem cells, which can promote recurrence and metastasis. Initially we hypothesized that obesity induced an activation of classical GC signaling which may promote stemness. However, known GR transcription targets *Nrcl1* and *Fkbp51* were not altered in the tumours following dexamethasone treatment (Fig. 2D). This indicates that dexamethasone use in obesity may promote stemness via a non-genomic signaling mechanism. This may involve either membrane or cytoplasmic GR that does not require nuclear translation and subsequent GR-mediated transcription. GR interacts with several kinases such as JNK, Src, PI3K, [19] and this may play an important role in the GC regulation of stemness, however, the non-genomic activities of GCs are not well understood. In summary, we provide pre-clinical evidence that administration of dexamethasone in obesity results in an increased cancer stem cell like subpopulation in mice and GCs may promote tumour reoccurrence in breast cancer patients also living with obesity

## Figure Legends

**Supplementary Table 1:**
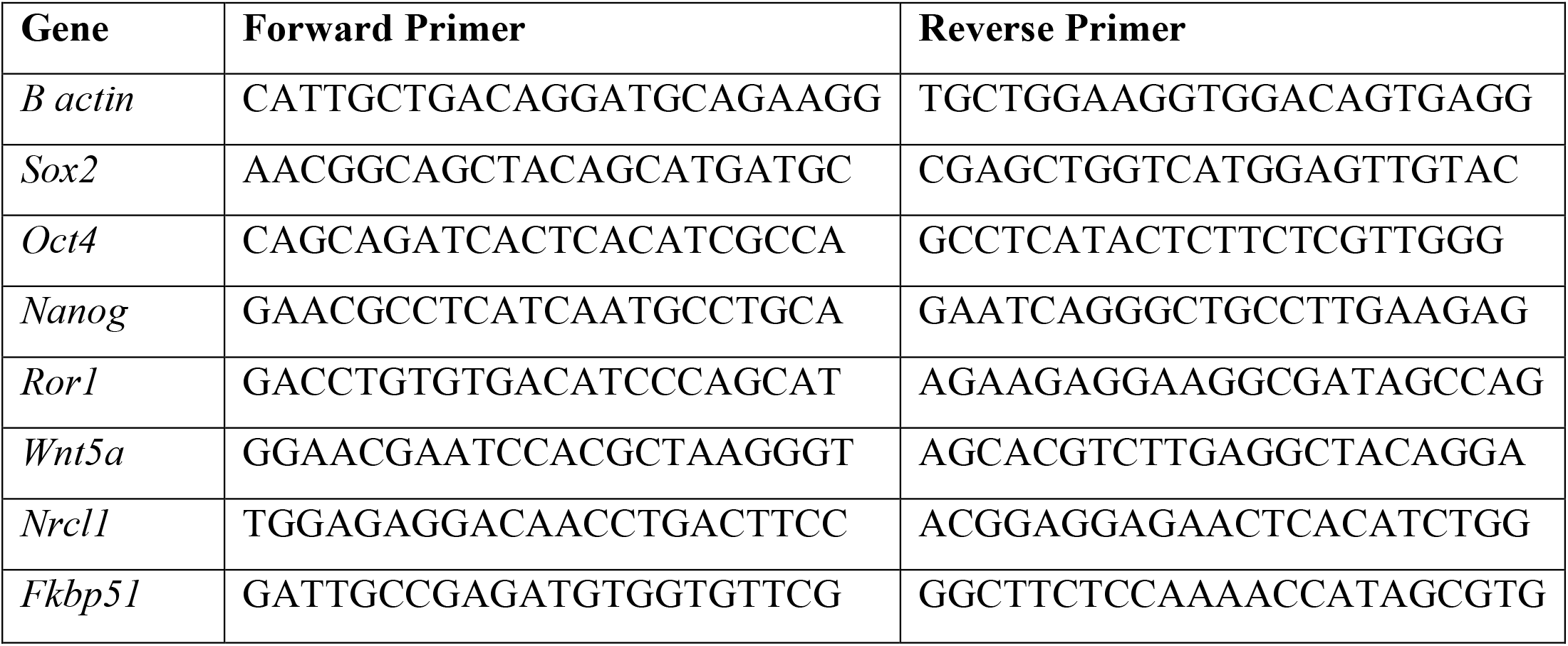
Real-time PCR primer sequences.

